# Computational approach and functional analysis of the *Pectobacterium carotovorum subsp. carotovorum* low-molecular weight bacteriocin Carocin S2

**DOI:** 10.1101/2020.08.19.257576

**Authors:** Jyun-Wei Wang, Ruchi Briam James S. Lagitnay, Reymund C. Derilo, Jian-Li Wu, Duen-Yau Chuang

## Abstract

*Pectobacterium carotovorum subsp. Carotovorum* 3F3 is a gram-negative phyto-parasitic enterobacterium. This strain is a producer of Carocin S2 bacteriocin, which comprises of two proteins of different sizes. Carocin S2K (killer protein) which is responsible for antibiotic resistance and Carocin S2I (immunity protein) which inhibits the antibiotic activity.

The present study aimed to predict the structure and functional properties of Carocin S2. Computational approaches utilizing various bioinformatic tools predicted that Carocin S2 is a putative membrane protein having the N-terminal at the extracellular side and the central domain at the coiled-coil region. Carocin S2 was predicted to have three domains, the translocation domains, receptor binding domain and the killer domain. Moreover, the killer domain was calculated to have the catalytic cleft. The in-vivo assays confirmed that for Carocin S2K, bound immunity protein was not a pre-requisite for cell attachment or translocation. The site-directed mutagenesis experiment led us to hypothesized the hydrolysis mechanism of Carocin S2.

The predicted structure of Carocin S2K provided a system of understanding on the biochemical and structural function which led to the mechanism of Carocin S2. It revealed that the role of immunity protein to Carocin S2 is not a pre-requisite for translocation pathway. Furthermore, this research led to hypothesized a hydrolytic mechanism of Carocin S2 to target the tRNA.

## Introduction

Bacteriocins are antimicrobial peptides produced by a bacterium that’s capable of inhibiting the growth of another microorganism. they’re classified as proteinaceous toxins, bactericidal, and are produced by both Gram-positive and Gram-negative bacteria (1). they vary from small lanthipeptides produced by lactic acid bacteria to much larger multi-domain proteins of gram-negative bacteria like the colicins from *E. coli*. Despite huge differences in chemical structures and post-translational processing of bacteriocins between the tiny (un)modified lanthipeptides produced by lactic acid bacteria and therefore the much larger multi-domain polypeptides produced by gram-negative bacteria like the colicins, all bacteriocins are ribosomally synthesized proteinaceous toxins that share a standard biosynthetic pathway. Bacteriocins are generally narrow-spectrum antibacterial agents with biological activity against closely related species and their genes are localized on transposable elements, plasmids, or on the chromosome of the producer’s genome (2). Bacteriocin production generally occurs following a stress signal, like DNA damage, inducing expression, and release of bacteriocin from the autolyzed cell (3).

Pectobacterium carotovorium subsp. carotovorium (Pcc) is a phytopathogenic enterobacterium accountable for the soft-rot disease of the many plant species. Some *Pectobacterium carotovorium* species produce one or more bacteriocin which boosts their competitiveness with other related bacterial species (4).

Previous studies revealed that Pcc produces Carocin S1, S2, and S3, each of which possessed different characteristics (5,6). They contain two proteins, one liable for the antimicrobial activity (the killing protein) and therefore the other for immunity (the immunity protein). The killing proteins are organized in functional domains with receptor binding, translocation, and DNAse or RNAse activity (7).

Proteins is easily understood in terms of its structure. Its three-dimensional structure is closely associated with its biological function. Proteins that perform similar functions tend to indicate a big degree of structural homology. amino acid sequence analysis provides important insight into the structure of proteins, which successively greatly facilitates the understanding of its biochemical and cellular function (8).

Several Bioinformatics tools are developed to predict the structure and functional properties of biomolecules (9). It provides a various array of computational tools for functional annotation and characterization of the expression of genome products. The computational genomics and proteomics analysis combined with predictive functional analysis represents another way for the rapid identification of latest putative bacteriocins similarly as new potential antimicrobial drugs (10).

This work aims to predict the structure and functional properties of the Carocin S2 from Pcc to the best possible accuracy vis a vis *in-vivo* study. Understanding these characteristics would enable us to deduce the hydrolytic mechanism, resulting in an improved understanding of the bacteriocin regulation mechanism.

## Results

### Computational analysis approach

An amino acid sequence was deduced from the nucleotide sequence of the *carocin S2* gene using DNASIS-Max software (Hitachi, Japan). The caroS2K was observed to be highly conserved among the different strains (*Pectobacterium carotovorum subsp. brasiliensis* PBR1692, *Serratia proteamaculans* (strain 568), *Pseudomonas aeruginosa* (strain PA7), *Escherichia coli* (ColD-CA23), *Pseudomonas aeruginosa* (strain LESB58), *Klebsiella oxytoca*,). Even though there were several variations in the nucleotide sequences as observed in the multiple sequence alignment of CaroS2K to other bacteriocins (Figure 1) it demonstrated good conservation at the C-terminal amino acid level. Multiple sequence alignment and phylogenetic analysis of both nucleotide sequences and protein sequences (UniprotKB_F8RN7, UniprotKB_P17998, UniprotKB_Q5QGN1, UniprotKB_A6V5R2, UniprotKB_Q06584) from different strains showed that they were identical.

**Figure 1.**
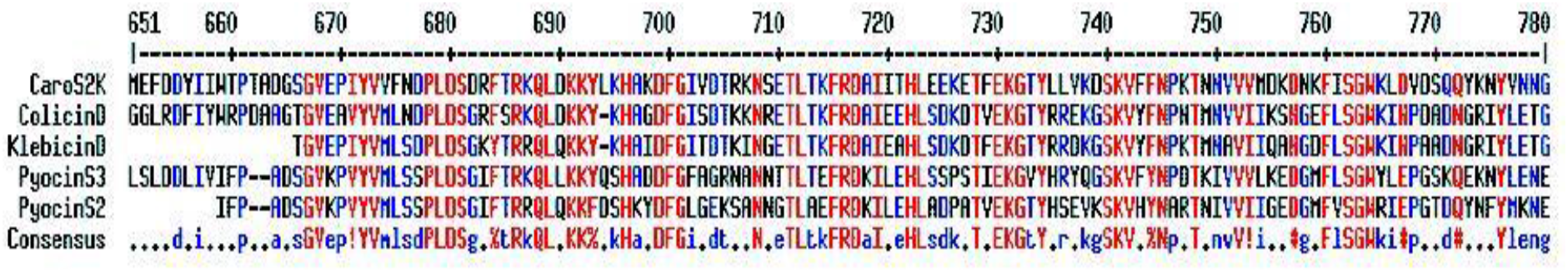
The multiple sequence alignment of CaroS2K with other bacteriocins.

Several prediction programs yielded widely discrepant results on the topology analysis of the CaroS2K sequence. Using the TMpred program, the predicted topology of the CaroS2K is IN-OUT with putative transmembrane segments (from amino acid positions 55-76, 93-115, 151-172, 373-392, 457-482, 470-499) (Figure 2). The N-terminal is predicted as present in the extracellular side of the membrane while the central domain is predicted as a coiled-coil region for receptor binding. Our consensus topology also passed the positive-inside rule and charge bias test, with a charge bias of+1 towards the inside of the membrane.

**Figure 2.**
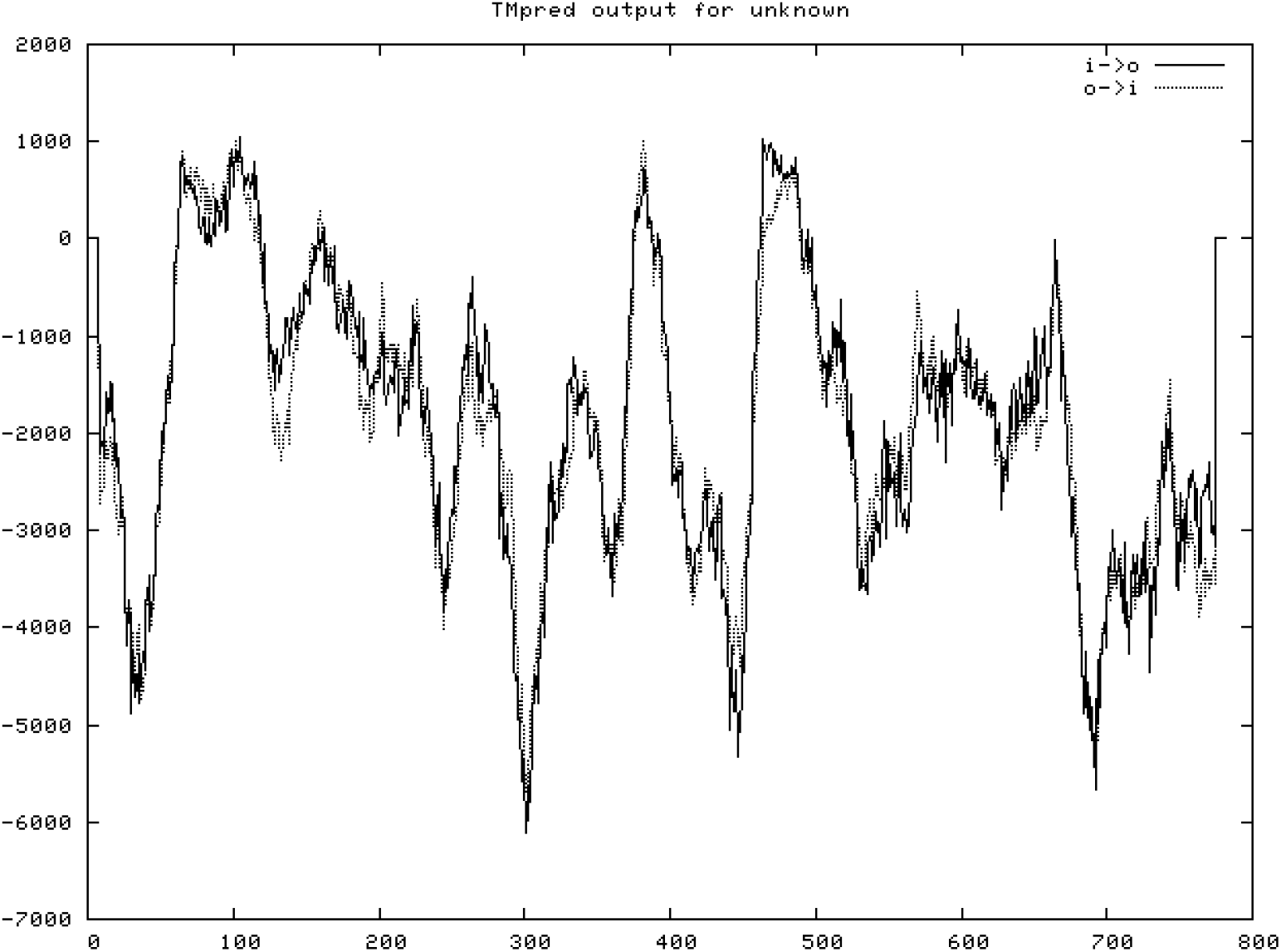
The topology prediction of Carocin S2K by TMpred program.

We sought the presence of domains, patterns, and motifs within the CaroS2K protein sequence to realize an insight into its functions and structure. The SMART results showed the presence of all the structural domains that we earlier identified using topology prediction programs. Our pattern search within the CaroS2K protein sequence, using different programs against the PROSITE database, gave similar results showing the presence of the many putative phosphorylation sites. All of the anticipated sites were found within the exposed regions of the CaroS2K.

Secondary structure identification of protein follows the three states, namely helix (H), extended (β-sheet) (E), and coil (C). However, to get a more consistent set of secondary structure, we’ve utilized the reduced version of DSSP (Database of Secondary Structure in Proteins) classification that uses eight kinds of secondary structure assignment: H (α-helix), E (extended β-strand), G (310 helix), I (π-helix), B (bridge, one residue β-strand), T (β-turn), S (bend), and C (coil).

In Figure 3A, Carocin S2K secondary structures were predicted by GOR, HNN and SOPMA protein programs. The ninety amino acid sequence showed that among the three programs, SOPMA predicted the highest alpha-helix structure (77.7%) while GOR predicted 43.33%. The extended strand was predicted 15.56% by GOR, 12.22% by HNN and 5.56% by SOPMA. Lastly, a random coil predicted by HNN tends to be 42.22%, while SOPMA predicted the least.

**Figure 3.**
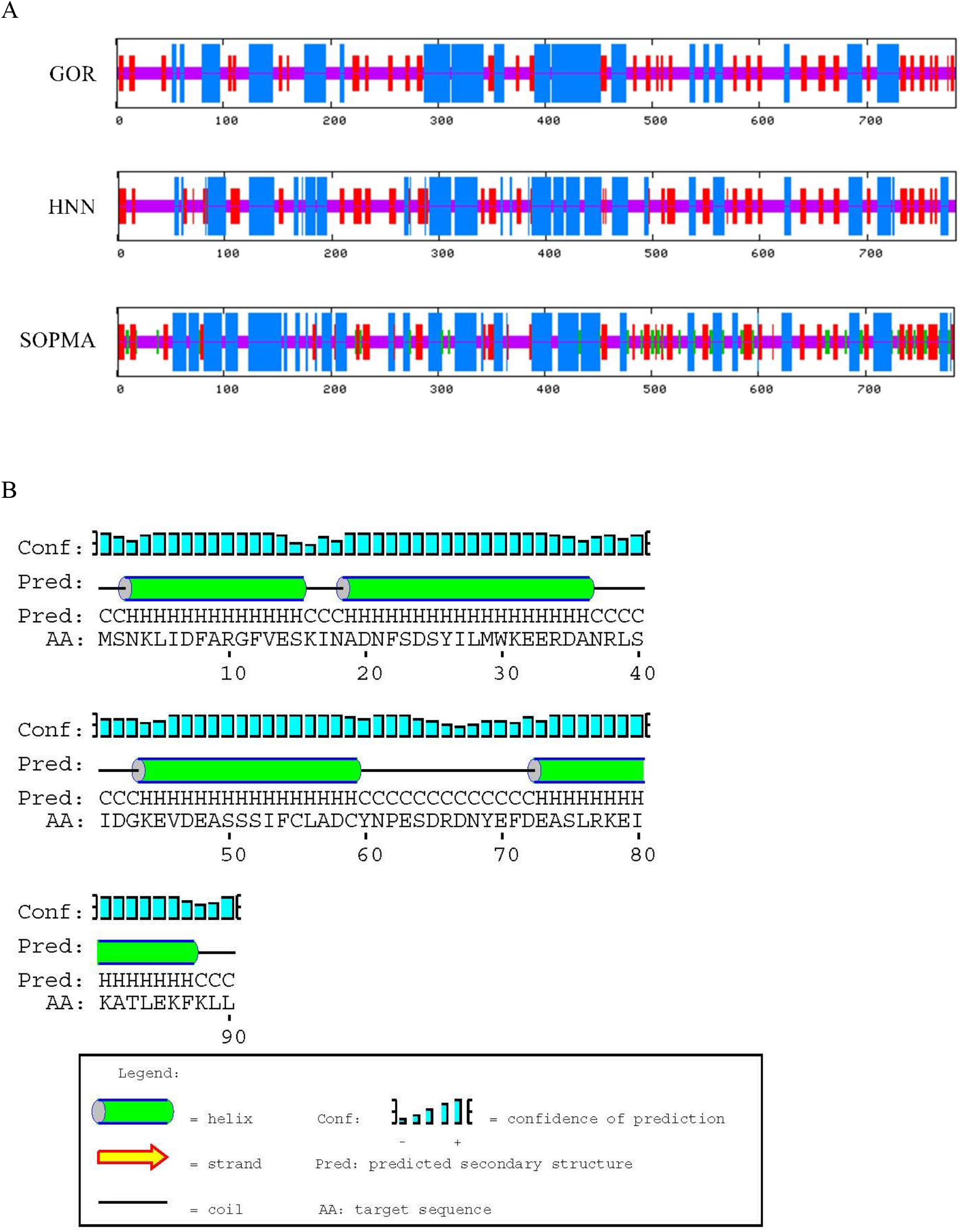
Predictions of CaroS2K structure. (A) The secondary structure elements prediction of CaroS2K by GOR, HNN, and SOPMA program. (B) The secondary structure prediction of CaroS2K by PSIpred program

For translation, we follow the strategy of CASP (Critical Assessment of Techniques for Protein Structure Prediction) and reduce the DSSP alphabet within the following manner: Helices (H, G, and I) within the DSSP code are assigned the letter H within the three-letter secondary structure code; whereas strands (E) and bridges (B) within the DSSP code are translated into sheets (E). Other elements of the DSSP structure (T, S, C) are translated into a coil (C).

PSIpred program was also utilized to predict the 9 amino acid sequence of Carocin S2K (Figure 3B). Diagrams annotate the query sequence with secondary structure cartoons and confidence value at each position within the alignment. Confidence is given as a series of blue bars.

Homology based tertiary structure prediction for the CaroS2K protein was present in Figure 4A, using Colicin D as the template.

The CaroS2K killer domain adopts a crescent-shaped globular α/β fold and consists of a central four-stranded antiparallel β sheet (β1, β2, β3 and β4) sandwiched between two long helices (α1 and α2) on one side and also the C-terminal helix (α3) on the opposite. As per the modeling prediction, the catalytic could be formed at the kinked helix (α1) lies within the plane of the β sheet, interacting with strand β4, parallel to β1*-loop-β2*.

**Figure 4.**
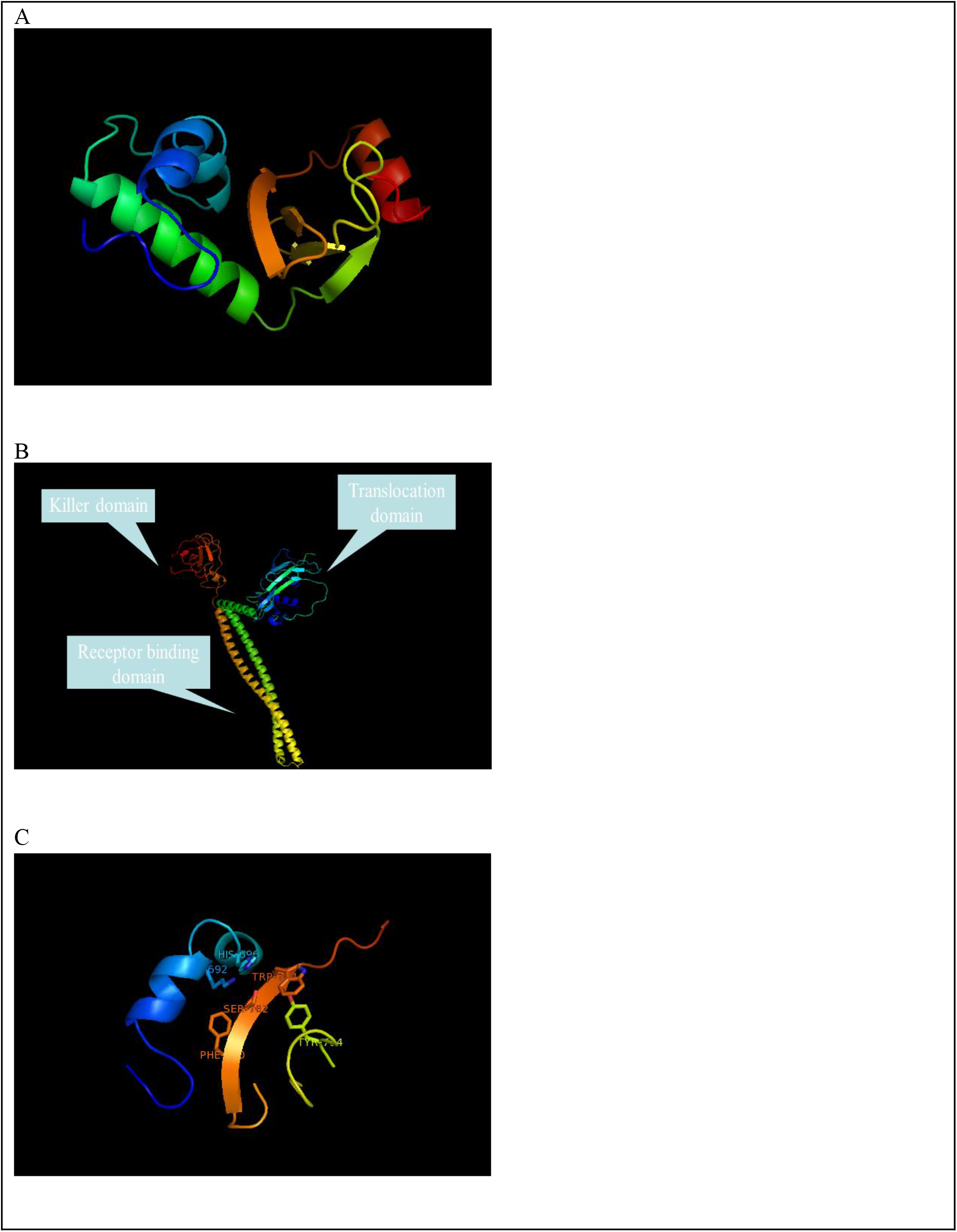
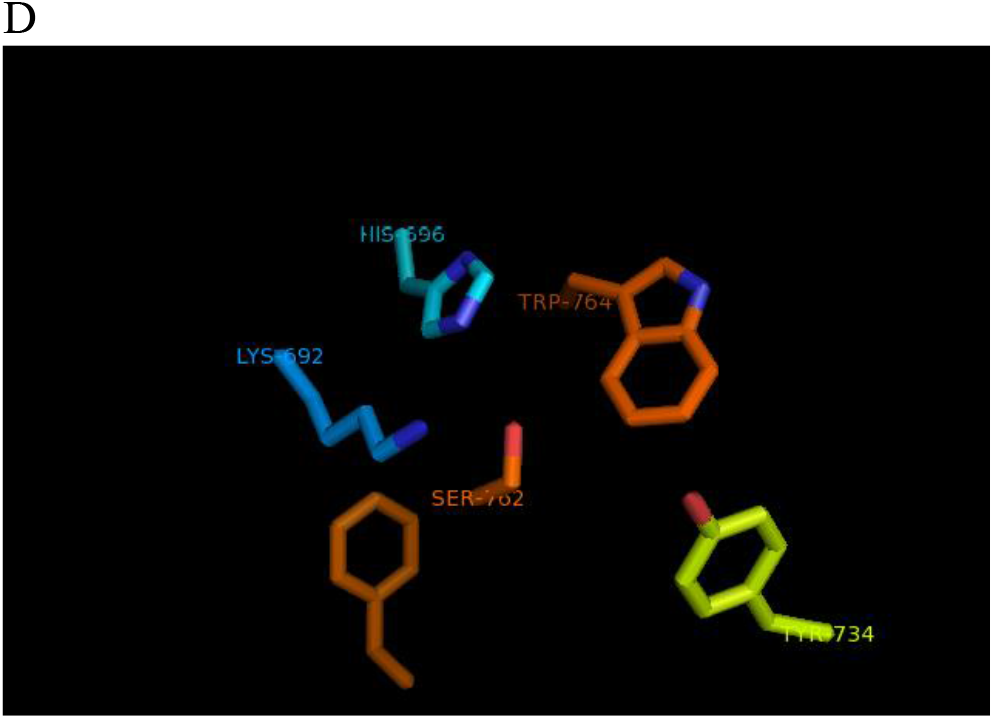
The CaroS2K modeling. (A) Homology modeling of CaroS2K by CPH models. (B) The threading method of CaroS2K by Phyre. (C) The catalytic clef of CaroS2K. (D) The functional residue of CaroS2K

We also used fold recognition-based structure prediction server Phyre to model the tertiary structure of the CaroS2K protein, using Colicin D, Colicin E3, and Colicin E7 as the template (Figure 4B). We refined the expected structure by fixing side chains, fixing problematic loops, removal of amino acid clashes (bumps) and energy minimization. The refinements didn’t yield any drastic change within the initial predicted structure. This was confirmed by visually inspecting the structure and verifying the backbone structure using the Ramachandran plot and computing the whole energy difference between the initial model and also the refined model.

The translocation domain formed a jelly-roll structure composed of the three β sheets flanked by two helical stretches (Figure 4B). The C-terminal element of secondary structure in the translocation domain was 30 Å long helix. At the end of this helix, there was a kink at Pro314, which led directly into the receptor-binding domain.

The receptor-binding domain consisted of 100 Å long antiparallel helical hairpins that extended from residue 315 to 455. The primary helical arm of this coiled-coil encompassed residues 315-380, with the last 3 residues of this stretch in an exceedingly 310-helical conformation. The second helical arm extended from residue 388-455. The antiparallel coiled-coil was “Ala coil” of the ROP type, during which every seventh residue was an alanine located within the core of the coiled-coil. The alanine residues occurred on alternate helices in every turn. There have been 3.5 residues per turn within the “Ala coil”, as distinct from a value of 3.6 for the canonical α helix. This was achieved by a smooth bending of the two helices around one another. There have been eight layers of alanine core residues within the receptor-binding domain. The receptor-binding domain was possibly the longest Alacoil ever observed. Such an antiparallel “Ala coil” is far tighter than the parallel leucine zipper.

The killer domain, consists of a four-stranded antiparallel β-sheet, with two helices on one side and a brief helix on the opposite side, as described above.

In the killer domain of Carocin S2, the triangle formation showed a similar arrangement, though these triangles are rather smaller than those of RNase A and T1. The bottom of the groove is rather hydrophobic (Phe760 and Trp764 from the β4 sheet). His696 is further surrounded by two polar residues: Ser762 (strand β4) and Tyr734 (β1*-loop-β2*) (Figure 4C and 4D). Helix α1 further carries three residues (Lys691, Lys692, and Lys695) creating a positive charge cluster that could play an important role for substrate binding by neutralizing the negatively charged phosphate groups of the substrate tRNA backbone.

### In vivo biological activity of CaroS2I

To confirm the activity of the predicted structure, in vivo biological activity of CaroS2I was done. The biological efficacy of overproduced CaroS2I was tested by establishing the level of immunity conferred to a sensitive strain of *Pectobacterium carotovorium* SP33. SP33 containing the expression vector pET30b was very sensitive to the bacterial activity of CaroS2I complex (Fig. 5A). This level of sensitivity was identical for SP33 without pET30b (data not shown). Constitutive expression of CaroS2I in SP33 resulted in complete resistance to CaroS2KI complex (Fig. 5B). Thus, expression of the gene for the CaroS2I protein conferred full biological immunity towards the action of its target Carocin, CaroS2K.

**Figure 5.**
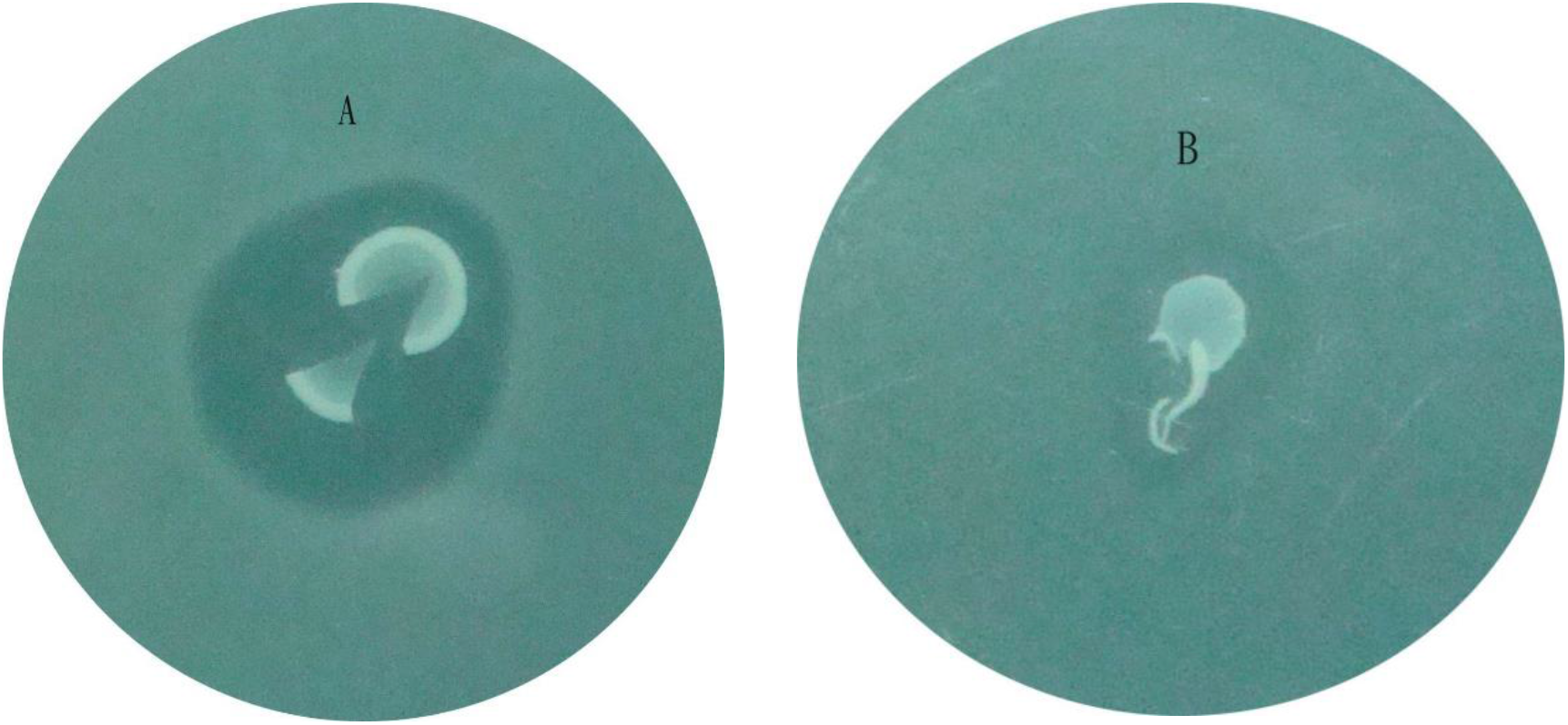
Overproduction of CaroS2I confers resistance towards the action of Carocin S2. A lawn of Pectobacterium carotovorium SP33 (A) containing the overexpression vector pET30b (without insert), and (B) the same strain and plasmid containing the CaroS2I. Zones, indicating cell death, were clearly apparent in the graph.

### Isolation and Characterization of unbound killer protein (CaroS2K)

Evidence from the electrophoresis from the previous study indicated that immunity protein may interact specifically with CaroS2K. Most purified Carocin S2 complex has a band that runs at the same position as purified immunity protein. Since the purification procedure for Carocin favors the separation of molecules with widely different ionic properties, it is unlikely that the two would co-purify to this extent if they were not interacting.

In figure 6A, it can be noted that spots 1, 2 and 3 demonstrated that Carocin S2K possessed a little activity against the indicator strain SP33 while spot 4 showed no activity.

**Fig. 6.**
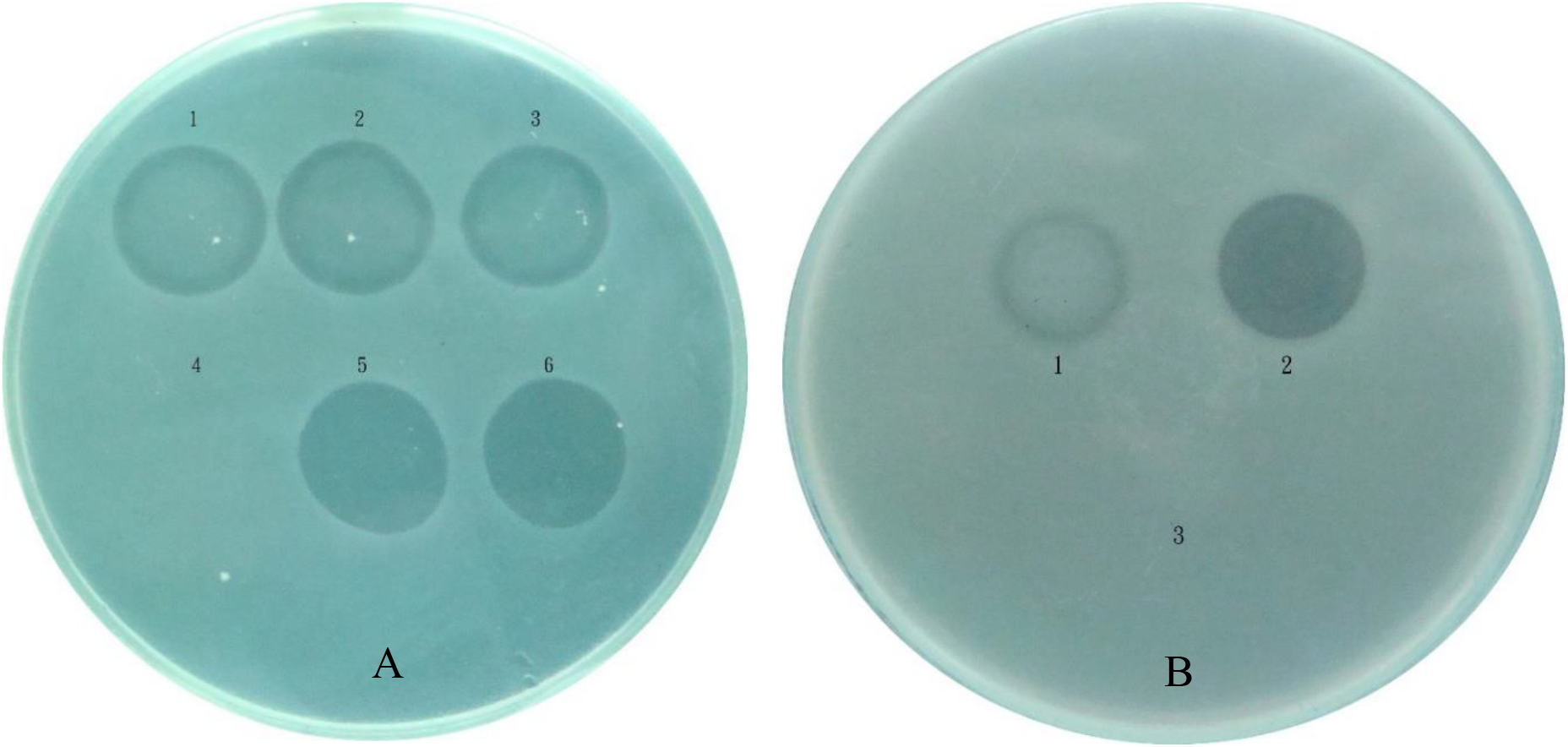
Inhibitory test of purified unbound killer protein (CaroS2K). (A) Clear zone of inhibitions are exhibited by; 1) crude extract of CaroS2K (2) purified CaroS2K under native condition (3) purified CaroS2K under denaturing condition with refolding buffer A (4) purified CaroS2K complexed to CaroS2I (5) purified CaroS2K under denaturing condition with refolding buffer B (6) purified CaroS2K under denaturing condition with refolding buffer C. (B) The inhibitory test of purified N2K, X2K, and C2K. (1) purified N2K (2) purified X2K (3) purified C2K

In general, the interaction between bacteriocin and its immunity protein is sufficiently strong that the separation can only be achieved under denaturing conditions. Denaturation by urea or guanidine hydrochloride was chosen as buffers to disrupt the interaction from Carocin S2KI-(His)6 complex. Moreover, spots 5 and 6 exhibited a clear zone of inhibition, which is a good indication that the unbound killer protein Carocin S2K was obtained successfully.

Furthermore, Carocin S2KI was also purified. The inhibitory activity of Carocin S2KI was shown in Figure 6B. Based on the figure, it may be observed that the expression of Carocin S2K without taq-protein at N-terminal (N2K) and Carocin S2K without taq-protein at C-terminal (C2K) has little or no activity against the indicator strain.

It has been shown that the Carocin S2K protein contains a catalytically active fragment capable of cleaving tRNA. Aside from Colicin E5, His residues play an important role within the catalytic mechanism of RNase. A minimum of one His, consistently present within the active site, is directly involved in proton abstraction of the 2’-OH ribose proton of the RNA substrate. Mutation of His696 abolishes its cleavage and toxicity activities. This His residue is found at the C-terminal end of the α1 helix, which forms the rim of the central groove of the protein. We next explored around His696 to find other candidate residues. Within the well-studied cases of RNase A and T1, the triangles formed by the catalytic triads (His12, Lys41, and His119 for RNaseA, and His40, Glu58, and His92 for RNase T1) display almost the identical geometric arrangement.

To clarify their functional involvement in tRNA hydrolysis, most of the residues mentioned above were turned into alanine and all of the mutants were compared to those of the wild-type bacteriocin-producing strain (Figure 7). The result revealed that Carocin molecules with mutated His696 or lysine residues forming the positive charge cluster (Lys692) are inactive. Mutant tRNAses affecting the most residues forming the central groove (Phe760, Ser762, and Trp764) or its rim (Tyr734) are not active. These structure-based mutagenesis experiments indicate that the central groove and also the positive charge cluster may form the active site of the Carocin S2.

**Fig. 7.**
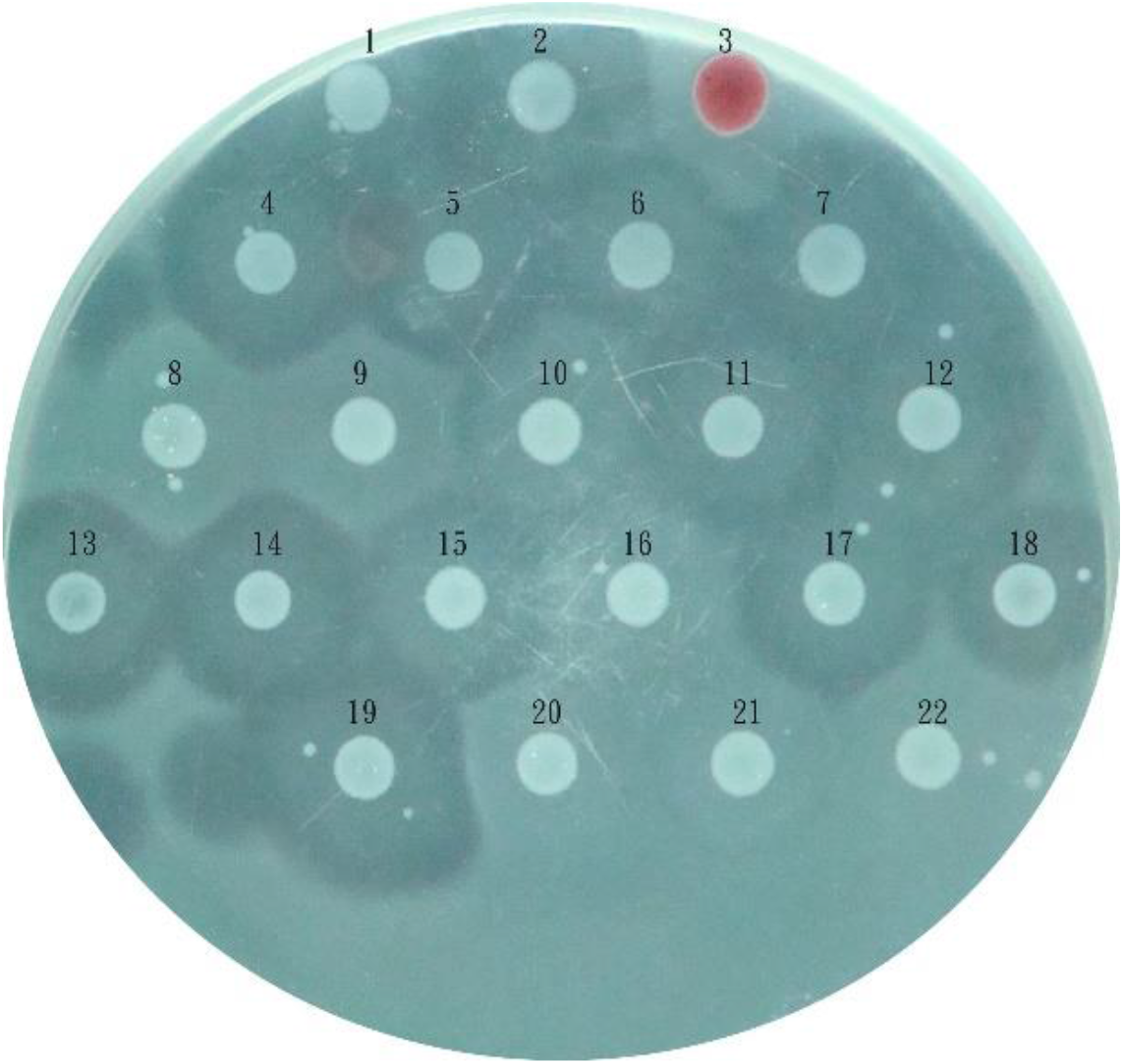
Mutation analysis of CaroS2K. (1) R320A (2) X2K (wild type) (3) Serratia (indicator) (4) T347A (5) K346A (6) T323A (7) K321A (8) H696A (9) K692A (10) K691A (11) K687A (12) T685A (13) D757A (14) D755A (15) S740A (16) Y734A (17) S709A (18) K698A (19) D767A (20) W764A (21) S762A (22) F760A

## Discussion

### Computational approach analysis for CaroS2K

Bioinformatics tools use a large type of algorithms to predict the properties of proteins at different levels. The accuracy of those Bioinformatics tools has been improving; however, each tool has its own advantages and downsides. A selected algorithm has its own characteristic specificity, sensitivity, robustness, computational cost, etc. These characteristics are often tested against benchmarks of known datasets (9). With the arrival of assorted sequencing techniques, amino acid sequences for a variety of proteins are determined. However, three-dimensional structural information obtained through X-ray crystallography, nuclear magnetic resonance, and other experimental methods are available just for around 10% of those protein sequences (10).

It was reported from the previous study that the sequence of Carocin S2 was compared to other analogous proteins using BLAST and FASTA search tools. It had been revealed that carocin S2 gene contains two open reading frames; one containing the 2352 bp caroS2K gene (translated to 85-kDa CaroS2K) and the other containing 273-bp caroS2I gene (translated to 10 kDa CaroS2I). The stop codon (TGA) of caroS2K overlaps the start codon of caroS2I by 5-bp (ATGA) (5).

Our prediction results showed that the CaroS2K has three probable domains – translocation domain, receptor binding domain, and killer domain. We also predicted that the CaroS2K contains many phosphorylation sites, ATP/GTP-binding site, and cAMP- and cGMP-dependent protein kinase phosphorylation site. The 3-D structure analysis of the CaroS2K also predicted the possible catalytic cleft within the killer domain. Thus, supported this computational analysis, and former experimental data, we hypothesized the hydrolytic mechanism of the CaroS2K. Additionally, CaroS2K contains only one cysteine residue per protein molecule. The relatively low proportions of hydrophobic amino acids and also the highly restricted number of cysteine residues are common properties of a variety of other extracellular proteins, including amylases, penicillinases, and flagella of varied species. These common properties of primary structure are also an element within the process by which the extracellular proteins pass through the cell membrane and cell wall.

### The role of immunity protein and unbound killer protein

Generally, bacteriocin operon consists of a killer gene, which encodes a killer protein that inhibits the growth of susceptible cell and an immunity gene, which encodes a protein which offers specific immunity against the killer protein (11,12).

Extracellular killer protein performs a biological action utilizing its receptor-binding domain which initially recognizes and connects the specific receptor on the membrane surface and further performs the mode of importation and also the translocation domain delivers the killer protein into the particular target within the susceptible cell (13).

In particular, the C terminal domain of killer protein determines the attacking mode while the killer protein gains entry into the infected cell. The bacteriocin producer is capable of specific immunity to the damage of its bacteriocin since the simultaneous production of the cognate immunity protein that ordinarily interacts with the C-terminal domain of the killer protein (14,15). It is notable that the immunity protein and the killer proteins interact at very high affinity due to charge attraction and are separated at the cell surface through the energy generated from the proton motif force (16).

This study represents the first of a series of investigations into the biochemical basis for immunity towards the RNase CaroS2Kfrom species of Pectobacterium. We have described here the purification and properties of the unbound killer protein and also the protein liable for immunity to CaroS2K.

Immunity protein was probably interacting with the CaroS2K. As stated within the previous study, the presence of purified Carocin S2 complex band that co-electrophoreses with immunity protein on SDS acrylamide gels strongly suggested that the two interacted Our results demonstrated clearly that, for CaroS2K, bound immunity protein was not a pre-requisite for cell attachment or translocation, since the free Carocin has precisely the same bacteriocidal activity (when analyzed by the zone spot test shown in Figure) as Carocin complexed to its immunity protein.

The in vivo activity of the overproduced CaroS2I was clearly observed by the whole protection it affords Pectobacterium carotovorium SP33, a strain which is generally sensitive towards the cytotoxic effects of CaroS2K. However, immunity protein incubated with CaroS2K so assayed in vitro for its protection of sensitive cells appears to be fully inactive. Therefore, whatever its mechanism of action, immunity protein cannot pass into Carocin-sensitive cells or perhaps prevent the attachment of the Carocin to its cell surface receptors. Otherwise, the immunity protein treated in vitro with CaroS2K would move in vivo. These properties are not expected for a molecule whose role was to shield Carocin-producing cells, but not the cells which are the targets of the Carocin.

### Hydrolysis mechanism prediction of CaroS2KI

The general catalytic mechanism by which ribonucleases hydrolyzed RNA has been thoroughly studied by protein engineering and crystallographic analysis over several decades. RNase A, for example, has two active sites His residues that cooperate during the catalytic cycle. One acts as a general base and abstracts a proton from the ribose 2’-OH, thereby catalyzing the nucleophilic attack of this radical on the 3’-phosphate group, resulting in a cyclic intermediate, while the opposite serves as catalytic acid during the first cyclization step. Their catalytic roles are reversed during the next hydrolysis of the cyclic intermediate. Other ribonucleases, like barnase and colicin E3, proceed probably through the same mechanism, but with a His and a Glu as catalytic residues.

We probed the potential active site region by systematic site-directed mutagenesis of residues on the catalytic surface of CaroS2K (Figure 8). Taken together, the results from the site-directed mutagenesis experiments show a clear picture of the catalytic pocket centered around His696. The substitution of this His by Ala ends up in the loss of the catalytic activity, supporting the expected structure that His696 is the general base of the cyclization step of the reaction. The nucleophilic attack of the 2’ -OH of the ribose on the 3’phosphate group creates a charged pentacovalent transition state that may be stabilized by nearby charged residues. In CaroS2K, the amine groups of Lys692 well-positioned to play such a task. Their mutation to Ala abolishes both cytotoxic and RNase activities of CaroS2K, confirming their role in tRNA substrate recognition or more directly within the catalytic activity. Mutation of Ser762, whose radical points towards His696, abolishes both cytotoxic and RNase activity, suggests a task for Ser762 in substrate binding and/or catalysis. Tyr734, Phe760, and Trp764 form a hydrophobic platform near the His696 imidazole side chain that exposes one face of its indole ring to the solvent, and will, therefore, stacks on a tRNA base. The lack of activity for the Tyr734Ala, Phe760Ala, and Trp764Ala mutant is in agreement with this proposal. Site-directed mutagenesis allowed us to map the CaroS2K active site and to spot residues critical for cytotoxicity (Lys692, His696, Tyr734, Phe760, Ser762 and Trp764).

**Fig. 8.**
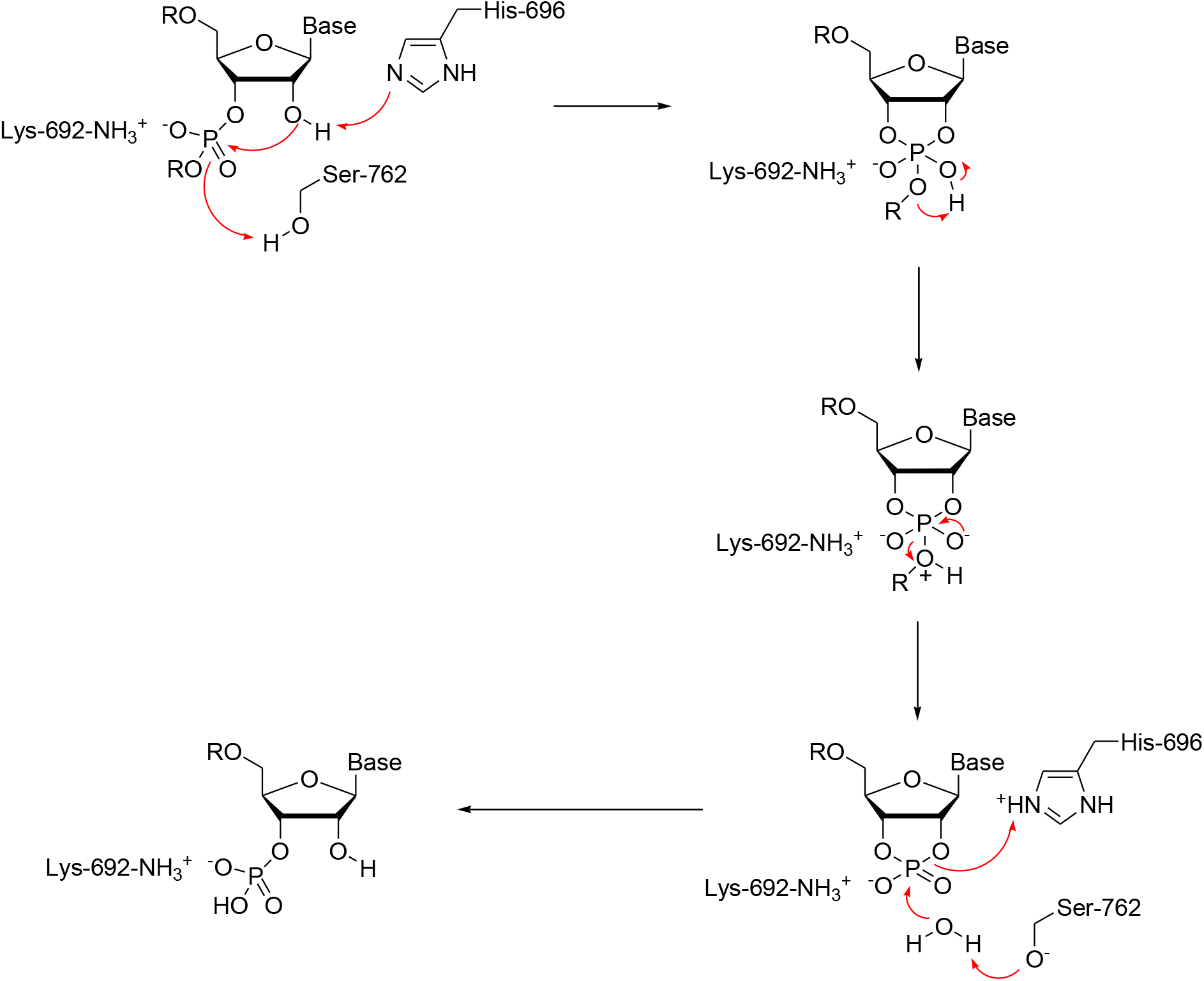
Hydrolysis mechanism prediction of CaroS2KI.

The substrate specificity of the tRNAse activity of CaroS2K is remarkably narrow. This activity was conferred by a small domain carrying a modestly sized active site cleft, and subtle recognition mechanisms must be at work to achieve this specificity. How this can result in a highly specific cleavage will have to await the structural information by crystallography.

### Experimental procedures

#### Bacterial strains and media

The bacterial strains, plasmids, and primers used in this study are listed in Table 1 and Table 2. Pcc strains were propagated at 28°C in a modified Luria-Bertani (LB) medium with 5g of NaCl per liter substituted for 10g. E. coli strains were grown in LB broth at 37°C with rotary agitation at 125 rpm. The antibiotic concentrations used for the selection of E. coli and Pcc strains were as follows: ampicillin, 50 μg/ml; kanamycin, 50 μg/ml; rifampicin, 50 μg/ml.

**Table 1.** Bacteria and Plasmids used in this study.

| Strain or Plasmid | Relevant characteristics | Source of reference |
| --- | --- | --- |
| ***Pectobacterium carotovorum* subsp. *carotovorum*** |  |  |
| F-rif-18 | Pcc, Rif<sup>r</sup>, wild-type | Laboratory stock |
| SP33 | Wild type, indicator | Laboratory stock |
| ***Escherichia coli*** |  |  |
| DH5α | supE44ΔlacU169(ψ80 lacZΔM15)hsdR17 recA1 endA1 gyrA96 thi-1 relA1 | Laboratory stock |
| BL21 (DE3) | hsdS gal(λcIts857 ind1 Sam7 nin5 lacUV5-T7 gene 1) | Laboratory stock |
| **Plasmid** |  |  |
| pGEM T-Easy | Amp<sup>r</sup>; lacZ cloning vector | Promega |
| pET30b | Kan<sup>r</sup>; expression vector with the C-terminal His-tag | Novagen |
| pET32a | Amp<sup>r</sup>; expression vector with the N-terminal His-tag | Novagen |
| pEN2K | *caroS2K* subcloned into pET32a | This study |
| pEX2K | Derived from pEN2K; deleted series of Tag-element in front of expressed *caroS2K* | This study |
| pEC2I | *CaroS2I* subcloned into pET30b | This study |
| pEX2I | Derived from pEC2I; deleted series of Tag-element in front of expressed *caroS2I* | This study |
| pEH2K | Derived from pEC2I; adding (His)<sub>6</sub>-Tag adjacent to *caroS2I* | This study |

Kan<sup>r</sup>: Kanamycin; Rif<sup>r</sup>: Rifampicin; Amp<sup>r</sup>: Ampicillin.

**Table 2.**
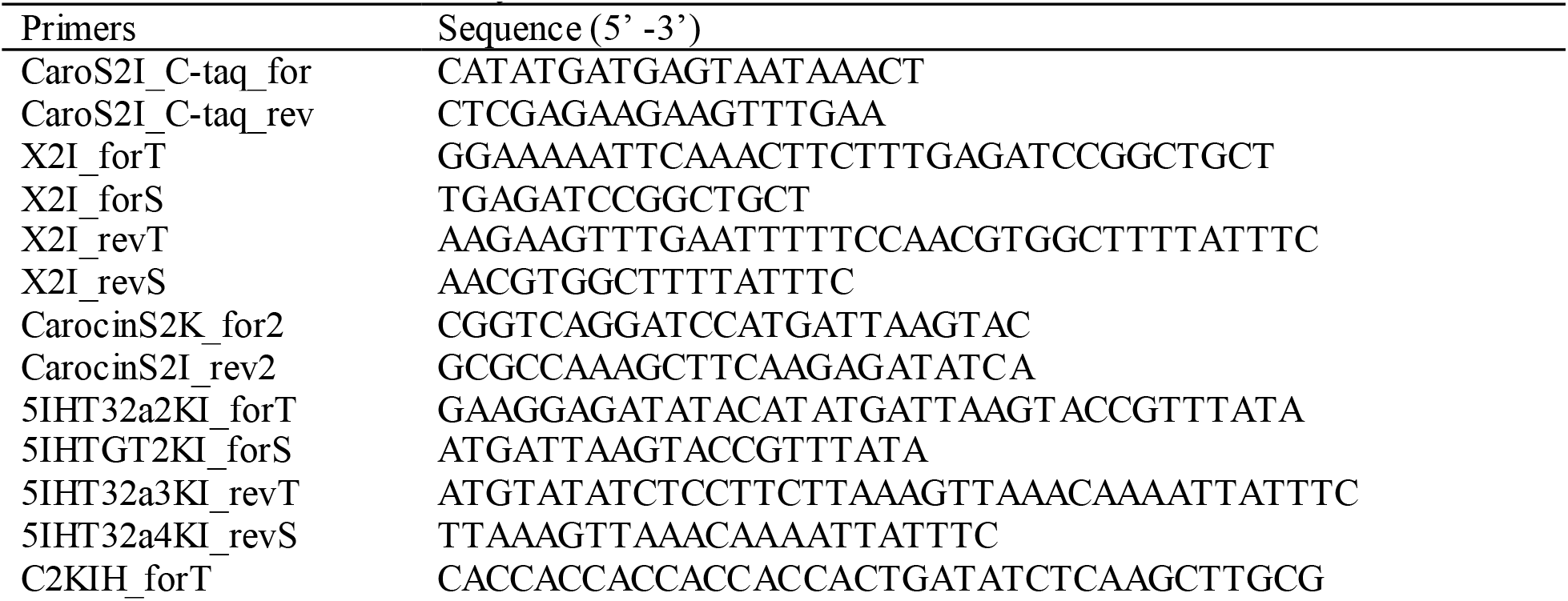
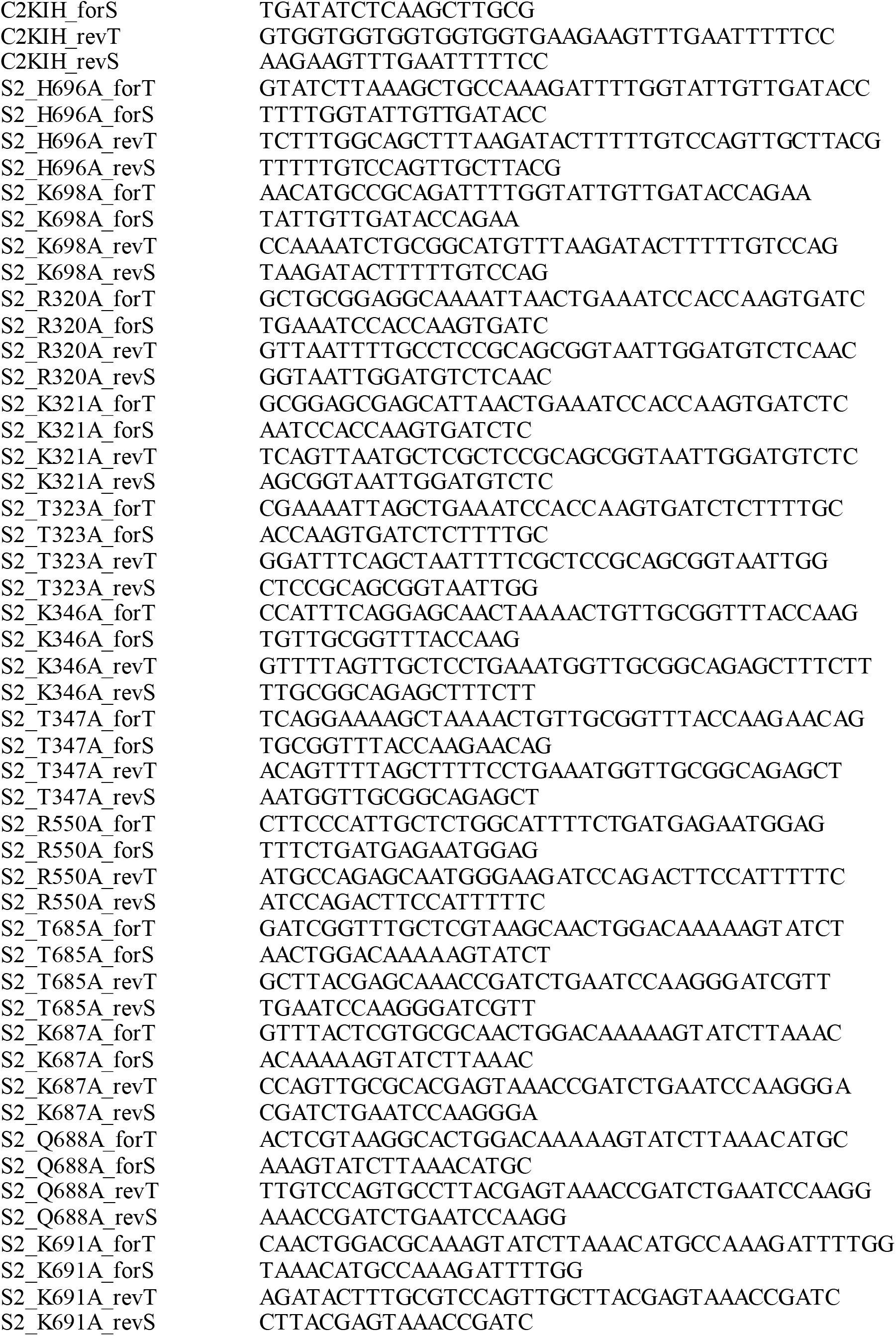
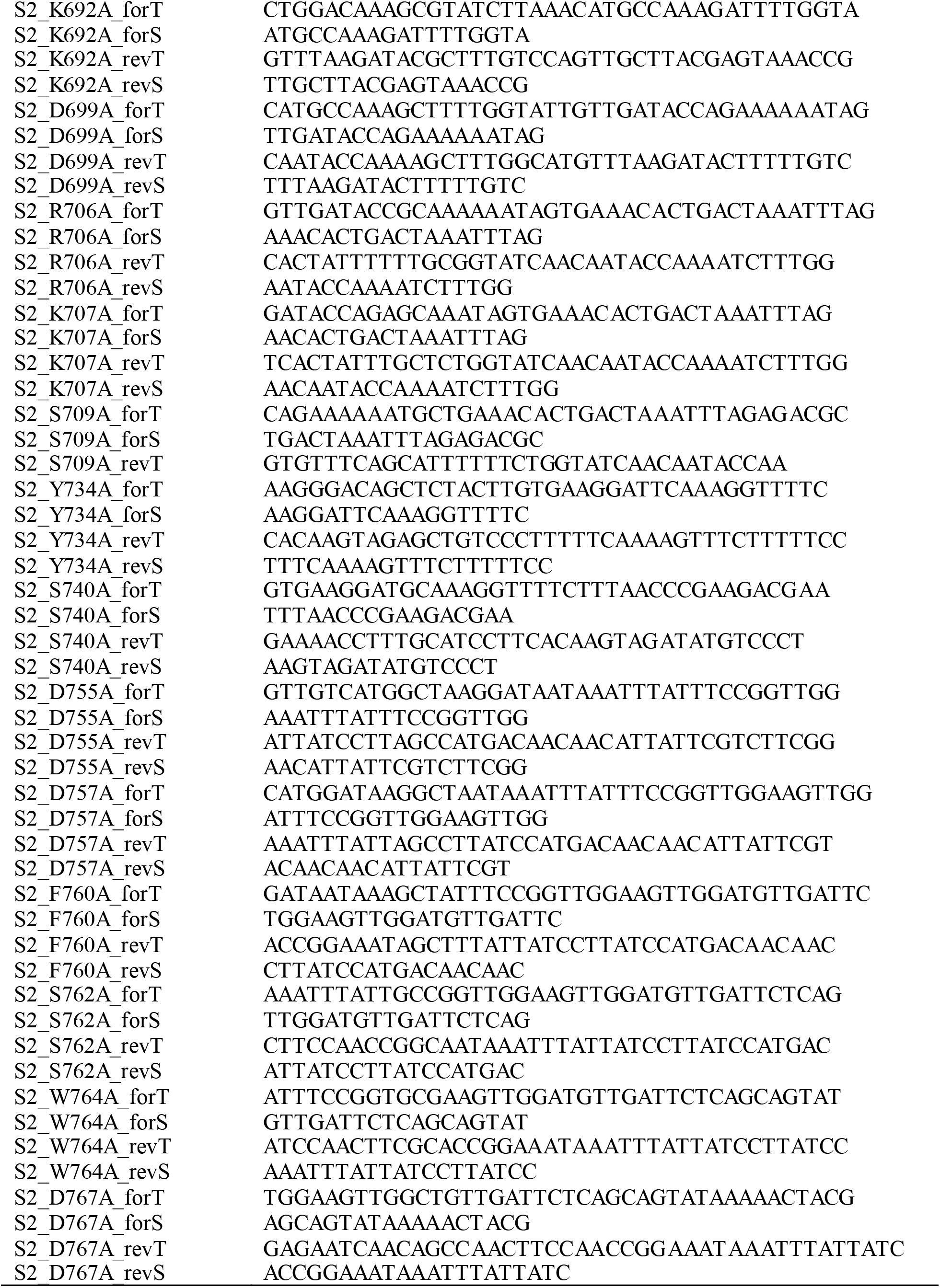
Primers used in this study.

#### Recombinant DNA techniques

Standard procedures for restriction endonuclease digestions, agarose gel electrophoresis, purification of DNA from agarose gels, DNA ligation, and other cloning related techniques were followed (17) with the use of the various primers listed in Table 2. The BLAST program was used to find the protein-coding regions.

#### Plasmid construction and DNA manipulation

The immunity protein of Carocin S2 was amplified from the genomic DNA of strain F-rif-18 with primers CaroS2I_C-taq_for and CaroS2I_C-taq_rev (Table 2). The construct pET30b-S2I (pEC2I) contains 273-bp amplicon was purified from the agarose gel, digested with *Nde*I and *Xho*I, and ligated onto *Nde*I - *Xho*I-linearized pET30b (Novagen). For further investigation, the CaroS2I without (His)_6_–taq protein was obtained by the method described previously with the following primers: X2I_forT, X2I_forS, X2I_revT, and X2I_revS (Table I). The pEX2I construct was then isolated from the kanamycin-resistant bacterial transformants, sequenced on both strands, and transformed into Escherichia coli BL21 (DE3) cells (Novagen).

#### PCR site-directed mutagenesis

A mutagenic primer was used in a PCR reaction together with either the universal primer or the reverse primer, whichever was appropriate. After isolation of the plasmid product with a Wizard Plus SV Minipreps (Promega), the PCR reaction was performed using the method described previously (18).

#### Protein expression and purification

Transformant of BL21, harboring protein expression from the plasmids pES2K or pES2I were grown in 500 mL LB medium (OD595 0.4). It was induced with 0.1 mM isopropyl-β-D-thiogalactopyranoside (IPTG) and were incubated at 25°C for 12 hours. Subsequently, the pellets were sonicated for 10 cycles for 9 seconds with 9-sec intervals. BL21/pES2KI pellet was harvested by ammonium sulfate precipitation (30%-40%) and was resuspended in buffer A (30 mM NaCl and 20 mM Tris-Cl; pH 8.0). The extract was equilibrated with buffer A and eluted with NaCl gradient (30 mM-1.4 M) was run with a fractogel column (Merck, USA). Purification was done with P-100 gel-filtration column (BioRad, USA) and the CaroS2K fraction were subjected for concentration using Amicon centriprep-50 column (Millipore, USA) and was dissolved in buffer A. BL21/pES2I pellets were precipitated by ammonium sulfate (70-100%), and were resuspended with buffer A. Similar purification process was followed for CaroS2I, however, it was concentrated using Amicon centriprep-3 column (Millipore, USA).

#### Antibacterial activity assay

The antibacterial activity of the bacteriocin producer strain was determined using the soft agar overlay method (19). Overnight cultures of the indicator strain Pcc SP33 were inoculated in prewarmed LB with an optical density of 0.5 at 600 nm, followed by aliquoting 100 μL into tubes with various Carocin concentrations. Sodium phosphate buffer (30 mM NaCl, 20 mM NaH2PO4 (pH 8.0) was used to prepare constant volumes for this inhibitory assay. Subsequently, the tubes harboring the indicator strains and Carocin were incubated with shaking at 28°C for one hour. The culture was plated on the LB plate and then incubated at 28°C for 16 hours. The production of an antibacterial substance was indicated by the presence of growth-free inhibition zones (clear zones) around the spotted area (20).

### Computational approaches to analysis

#### Primary sequence analysis

The protein sequence was obtained by conceptual translation of the caroS2 open reading frame from the P.c.c genome. The primary sequence analysis was performed using ProtParam and ProtScale (21). ProtScale was used to predict the CaroS2 profile based on several amino acid scales. ProtParam computed various properties like molecular weight, theoretical pI, instability index and grand average of hydropathicity.

#### Similarity searching

Different programs were used from NCBI like BLASTP and PSI-BLAST (22) and compared against several different databases like SWISSPROT and PDB. Multiple sequence alignment and phylogenetic analysis were done using Florence Corpet.

#### Topology prediction

The Topology of CaroS2 protein was predicted using TMpred (23), TMHMM (24), DAS (25), Coils, and MultiCoil. TMpred algorithm was based on the statistical analysis of TMbase, a database of naturally occurring transmembrane proteins, using a combination of several weight-matrices for scoring. TMHMM predicts transmembrane regions based on the hidden Markov model. DAS uses dense alignment surface method to predict transmembrane regions.

#### Secondary structure prediction

Programs GOR, HNN, SOPMA, and PSIpred in the NPS (Network Protein Sequence Analysis) consensus secondary structure server (26). This server runs the input sequence against several different secondary structure prediction tools and generates a consensus secondary structure out of them.

#### Domains/patterns/motif prediction

SMART (Simple Modular Architecture Research Tool) was used to identify the presence of any domains in the CaroS2 protein. Different pattern searching applications (ELM, Motif Scan, and PROSITE scan (27) were utilized to predict functionally relevant patterns in CaroS2 protein.

#### 3-D Structure prediction and analysis

Initial attempts to predict the tertiary structure of CaroS2 were done using different approaches like homology modeling and threading. Automated homology modeling servers, Swiss-Model and different programs (CPHmodels, ESyPred3D, and 3DJIGSAW) were used for homology modeling. Predictions described in this study were done using fold recognition tools LOOPP, and a new version of 3-D PSSM (Phyre). The quality of the predicted structure was examined using an online version of the WHATIF program. Structure refinement was done using both WHATIF and Swiss-PDB Viewer. Structure visualization was done using PyMOL, RasWin and VMD. The figures were produced using a combination of the programs’ PyMOL.

## Data availability

The datasets used and analyzed during the current study are available from the corresponding authors on reasonable request.

## Acknowledgements

Not applicable

## Author contributions

JW, RBJSL, RCD and JLW participated in the discovery and characterization of Carocin S2 and wrote the manuscript. DYC conceived the study, participated in its design and corrected the manuscript. All authors read and approved the final version of the manuscript.

## Funding and additional information

The support of this work by grants from Chang Bing Show Chwan Memorial Hospital, Taiwan, is gratefully acknowledged.

## Abbreviations and nomenclature

The following abbreviations were used in this study:

PCC: *Pectobacterium carotovorum subsp. carotovorum*
SMART: Simple Modular Architechture Research Tools
DSSP: Database of Screening Structure in Proteins
GOR: Garnier-Osguthorpe-Robson
HNN: Hierarchical Neural Network
SOPMA: Self-Optimized Prediction Method with Alignment
CASP: Critical Assessment of Techniques for Protein Structure Prediction

